# Genetic heterogeneity induces nonadditive behavioural changes in *Drosophila*

**DOI:** 10.1101/2024.08.19.608728

**Authors:** Takahira Okuyama, Daiki X. Sato, Yuma Takahashi

## Abstract

The formation and dynamics of group behaviours are important topics in ecology and evolution. Although several theoretical studies assume homogeneity among individuals, real-world organisms often display remarkable behavioural diversity within groups. This study investigated the synergistic impact of genetic heterogeneity on group behaviour and reveals the behavioural underpinnings of diversity effects using 83 genetically distinct strains of *Drosophila melanogaster*. Various indices of foraging behaviour, including movement speed, search comprehensiveness, spatial preference and stopping time, were measured using homogeneous (single strain) and heterogeneous (mixing two distinct strains) groups of flies. The heterogeneous groups exhibited significant increases in spatial preference and stopping time compared with the homogeneous groups, suggesting that genetic heterogeneity induces nonadditive changes in group behaviour. Furthermore, the magnitude and direction of the behavioural change varied among different combinations. Multiple regression analysis showed that the phenotypic distance in some traits between mixed strains could explain the emergence of diversity effects on group behaviour. Specifically, interindividual heterogeneity in the locomotor activity level showed a positive correlation with diversity effects. These results emphasise the importance of intraspecific diversity in group dynamics and suggest that genetic heterogeneity can improve group performance through the acquisition of novel behavioural traits.

## Introduction

Group living is a widespread phenomenon in the animal kingdom, and its emergence and evolution are important topics in ecology and evolution (Westneat et al., 2010). Groups of animals engage in behaviours as diverse as predator detection (Hamilton, 1971; Treherne and Foster, 1981), navigation (Biro et al., 2006; Weimerskirch et al., 2001) and social foraging (Barnard and Sibly, 1981; Giraldeau et al., 1999). Group formation can occasionally impose various costs on individuals within the group. A typical example is interference over resources, which is insignificant at low competitor densities but steadily increases in intensity as the density rises (Triplet et al., 1999).

Furthermore, aggregation can facilitate the transmission of pathogens or amplify the effects of stress (Brandl et al., 2022). Theoretically, animals form groups when the benefits of aggregation exceed the cost of maintaining close proximity with conspecifics (Markham et al., 2015). Hence, for understanding the dynamics of group formation, it is necessary to elucidate the mechanism by which individual benefits occur. Studies have suggested that group formation exerts various beneficial effects on individuals in groups such as reducing predation risk and improving foraging efficiency, wherein the former phenomenon is attributed to an increased probability of detecting predators with the heightened surveillance (Krause et al., 2002) and the reduction of predation pressure through dilution and confusion effects (Hamilton, 1971; Neill and Cullen, 1974). It is known that larger flocks of dove (*Columba palumbus*) take flight and avoid predators even when they are at a considerable distance (Kenward, 1978). The latter phenomenon, viz., improving foraging efficiency, is attributed to prey discovery through information sharing and interactions among individuals. For instance, the crow (*Corvus corax*) after discovering a food source actively communicates to other crows and leads the foraging group to the site (Marzluff et al., 1996).

The dynamic changes in collective phenomena can be seen as a consequence of local interaction among individuals (Reynolds, 1987). The behavioural traits of other members can influence one’s own behaviour, a phenomenon known as the indirect genetic effect. The two-spotted cricket (*Gryllus bimaculatus*) with aggressive genotype was shown to strongly decrease aggressiveness in opponents (Santostefano et al., 2017). Such social interactions can ultimately impact the dynamics of the group. In this respect, recent studies have paid much attention to the effects of interindividual heterogeneity on group behaviour (Jolles et al., 2020). For instance, the heterogeneity in boldness among individuals of guppies leads the synergistic behavioral changes and allows the group to explore a wider range of food resources, benefitting the entire population (Dyer et al., 2009). In *Drosophila melanogaster*, a population with behavioural diversity in locomotor activity has a higher survival rate than a population without diversity (Takahashi et al., 2018). Nevertheless, the effects of diversity on groups do not consistently manifest in the same pattern. For instance, in *Stegodyphus dumicola*, a species of spiders, a higher proportion of bold individuals in the population results in a higher infection rate by microorganisms (Keiser et al., 2018). These facts suggest that the effects of diversity on groups can be either positive or negative depending on the situation. However, it remains unclear which types of diversity lead to specific synergistic behavioural changes, including indirect genetic effect. The relationship between the type or strength of heterogeneity and its impact on groups dynamics is not well understood.

Therefore, we conducted this study to investigate the synergistic effects of intraspecific behavioural diversity on group behaviour using 83 genetically distinct strains of *D. melanogaster*. For quantifying the effects of intraspecific diversity on group behaviour, we established homogeneous (consisting of a single strain) and heterogeneous (consisting of two distinct strains) groups and recorded the behaviour of individuals in both groups. Next, we measured the moving speed, search comprehensiveness, spatial preference and stopping time as indices of food-seeking behaviour. We also explored the behavioural features of the combinations that exhibited the diversity effect to identify the type and strength of heterogeneity leading to synergistic behavioural changes in groups. Our study could provide novel insights into how intraspecific diversity, previously overlooked, can affect behavioural group characteristics.

## Materials & Methods

### Fly strains and rearing conditions

A total of 83 strains of *D. melanogaster* were obtained from the *Drosophila* genetic reference panel (DGRP), which consists of inbred lines derived from a single natural population in the United States (Mackay et al., 2012). Flies were reared for successive generations in a constant environment (12L:12D, 25°C) with lights on at 6:00 and off at 18:00. Adult females aged 0–2 days after emergence were collected and reared for 2 days in the same environment. During this period, 5 males and 15 females were placed in the same vial to ensure mating before being used in the experiment to prevent behavioural differences due to mating experience. Adult females aged 2–4 days were used for the behavioural assay.

### Behavioural assay and tracking

Behavioural assays were performed during a Zeitgeber time of 0–12 in an incubator maintaining the same abovementioned temperature conditions. Females were anesthetized with CO_2_, which is considered to exert relatively little impact on their behaviour (Perron et al., 1972), although the negative side effects of CO_2_ anaesthesia have also been reported (Colinet and Renault, 2012), and then introduced into the experimental arena with a diameter of 30 mm and a depth of 2 mm, where the flies were restricted from flying (Fig. 1a). Six females derived from either a single strain (homogeneous condition) or two strains (heterogeneous condition; three flies from each strain) were introduced into the arena. After 30 min of acclimation in the incubator, 12 arenas arranged on a single acrylic plate were set on an LED board (147 mm width × 115 mm height, 7500 lx white light, TLB-MP, Asone Co., Japan). After another 10 min of acclimation, their behaviour was filmed for 5 min. The experiments were conducted three to six times per strain or the combination of strains, and the position of the arena in a plate was randomly changed among replications.

**Figure 1.**
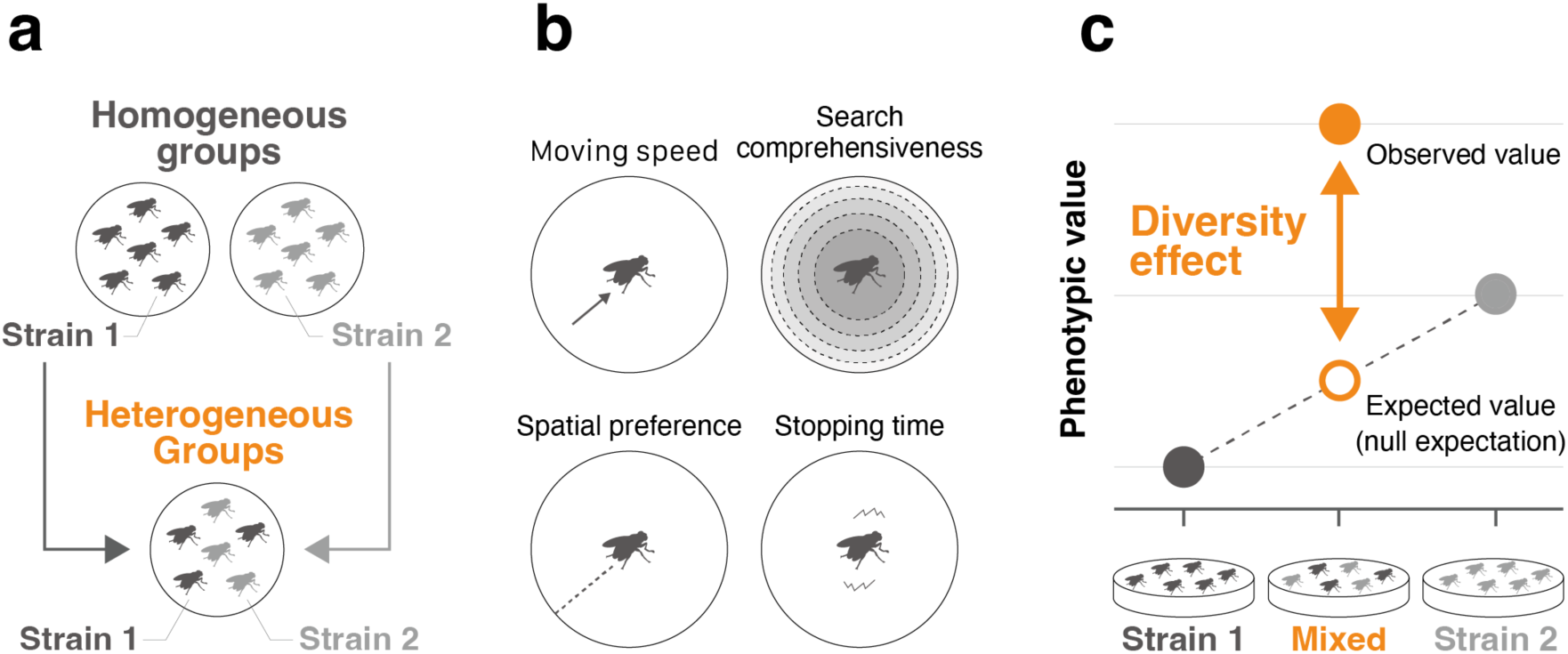
Experimental scheme. (a) We established a heterogeneous group by mixing two distinct strains. (b) Four behavioural indices related to foraging were quantified from the tracking data. (c) Diversity effects were calculated by comparing the observed value with the expected value.

The locomotive behaviour of flies was recorded at 1920 width × 1080 height in pixel resolution and approximately 50 frames per second using a USB3 camera (DMK33UX290, The Imaging Source, Germany) controlled by a custom Python script. The recorded videos were cropped for each arena, and the behaviour of flies was tracked using Flytracker (Eyjolfsdottir et al., 2014)to obtain data on their movement tracks.

### Indices of foraging and vigilant behaviour

We quantified the following four behavioural indices as foraging behaviour: moving speed, search comprehensiveness, spatial preference and stopping time (Fig. 1b). Tracked coordinates were standardised based on the position of the arena, i.e. the centre coordinate of the arena was set to (0,0), and the diameter was set to 30. We next calculated the distance travelled from a previous time point and the speed (travelled distance divided by frame intervals for each time point) with corrected coordinates and summarised them at 0.5-s intervals. Moving speed was calculated as the average speed of flies moving at speeds >4 mm/s. Search comprehensiveness was measured as uniformity of the explored area. The arena was divided into five concentric parts of equal area, and the Simpson’s diversity index (Simpson, 1949) of time spent in each area was calculated. Spatial preference was evaluated as a mean distance from the edge of the arena during observation. Stopping time was measured as the sum of the time when the flies moved <0.5 mm/s.

### Calculation of the diversity effect

We obtained 71 heterogeneous groups from the 83 distinct strains of *D. melanogaster*. To examine the impact of interindividual diversity on group behaviour, we compared the observed behavioural indices from the heterogeneous groups with the expected values from the homogeneous groups. This allowed us to identify the diversity effect, i.e. the nonadditive effects of interindividual diversity on group performance. For each trait of interest, the diversity effect was defined as the discrepancy between the observed behavioural indices in the heterogeneous groups and their expected values calculated as the mean of two homogeneous groups (Fig. 1c), as follows:

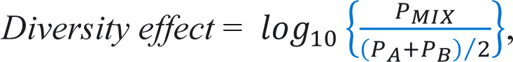

where *P_A_* and *P_B_* represent the phenotypic values of the homogeneous groups constituting a heterogeneous group, and *P_MIX_* represents the phenotypic value of the heterogeneous group.

For calculating the mean phenotypic value of each strain and each combination (i.e. *P_A_*, *P_B_* and *P_MIX_*), we used a linear mixed model, including each behavioural trait as an explanatory variable, strain as a fixed effect and plate, arena, experimental date and time of the day as random effects.

### Behavioural basis of diversity effect

The relationship between group composition and the emergence of diversity effect was clarified using multiple regression analysis, where the phenotypic distance for each combination was used as an explanatory variable, and the diversity effect for each combination was used as a response variable. Because each of the four measured behavioural traits correlated partially, they were summarised using principal component analysis (PCA). Then, the difference in each PC score between mixed strains was calculated as the phenotypic distance between strains. A squared term was also included in the model to account for parabolic correlations. To identify important variables for the response variable, the best-fitting model for multiple regression analysis was selected based on the AIC criterion.

### Statistical analyses

All statistical analyses were conducted using R 4.3.1. The Wald test was applied with the ‘car’ package for the analysis of variance to examine the variation among strains and to determine the statistical significance of each variable in multiple regression models. The ‘ggeffects’ package was used to predict the mean phenotypic value of each strain or combination by considering random effects. The best-fitting model was selected in the multiple regression analysis using the ‘MASS’ package. In this model, the number of replications for each strain or combination was used as a weighting variable to account for the varying sample sizes. Throughout all statistical analyses, standard errors were calculated to represent variation.

## Results

### Genetically diverse strains exhibit significant variation in behaviour

We first investigated behavioural correlations using the dataset of homogeneous groups from the 83 strains. Moving speed (χ^2^ = 919.4, *df* = 82, *P* < 0.001), search comprehensiveness (χ^2^ = 431.5, *df* = 82, *P* < 0.001), spatial preference (χ^2^ = 329.1, *df* = 82, *P* < 0.001) and stopping time (χ^2^ = 856.4, *df* = 82, *P* < 0.001) significantly varied among strains (Fig. S1). Correlations among the behavioural indices were significant, except for that between spatial preference and search comprehensiveness (Fig. 2a). The 71 heterogeneous groups also demonstrated significant variation in moving speed (χ^2^ = 496.5, *df* = 70, *P* < 0.001), search comprehensiveness (χ^2^ = 255.9, *df* = 70, *P* < 0.001), spatial preference (χ^2^ = 188.8, *df* = 70, *P* < 0.001) and stopping time (χ^2^ = 548.6, *df* = 70, *P* < 0.001) (Fig. S2). The phenotypic distance between two strains paired varied among the 71 heterogeneous groups. The variation was not uniform across the four behavioural indices, ensuring behavioural diversity within the heterogeneous groups (Fig. 2b).

**Figure 2.**
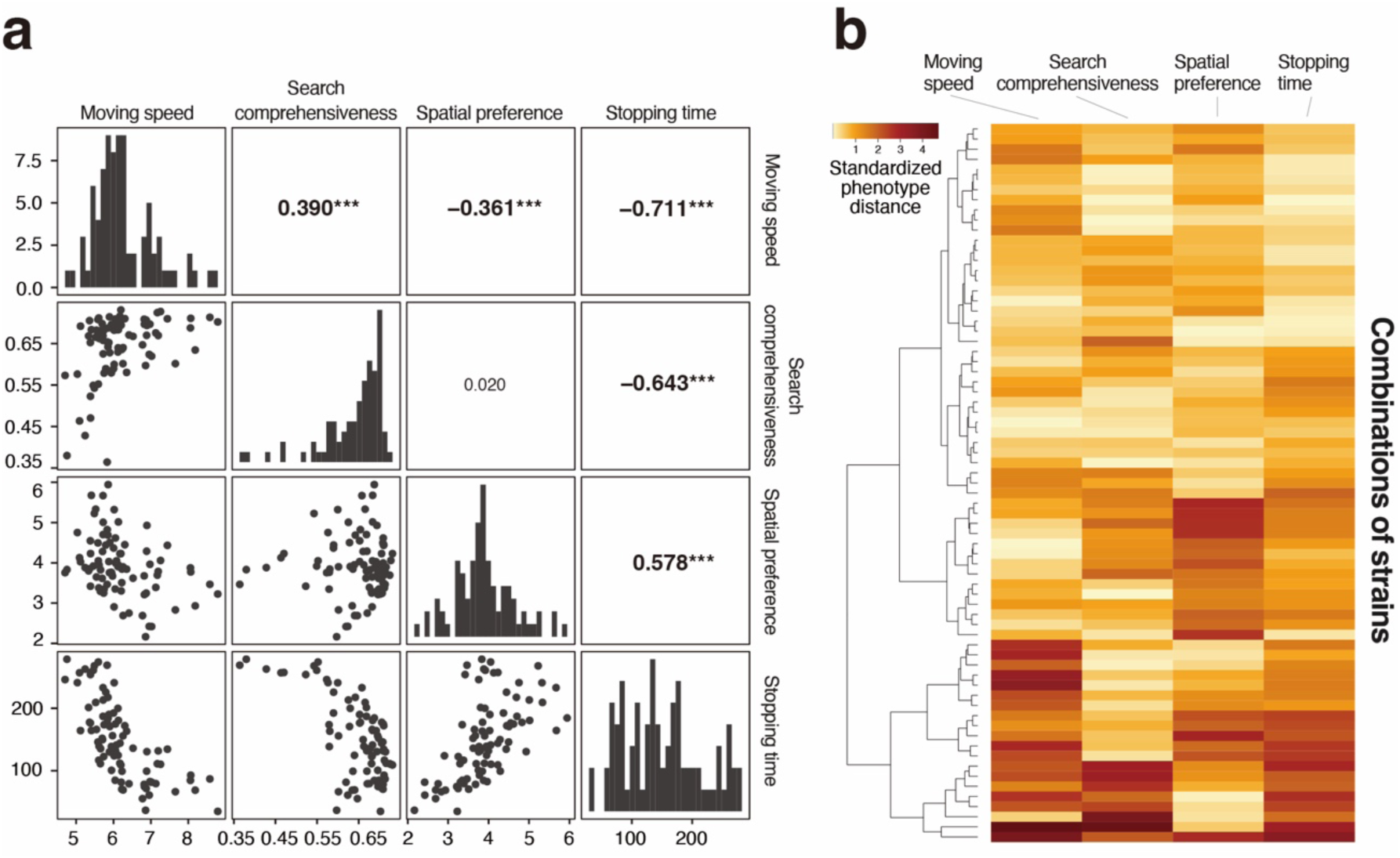
Correlation and variation in behaviour observed in genetically diverse fly groups. (a) Pairwise correlations among the four behavioural indices were examined using the dataset of homogeneous groups from 83 strains. Numerical values indicate Pearson’s correlation coefficients, and asterisks denote *P* values (*** *P* < 0.001, ** 0.001 ≤ *P* < 0.05, *0.001 ≤ *P* < 0.05). (b) For combinations of strains used in the heterogeneous group experiment, phenotypic distances were visualised in a heat map, with darker colour representing larger distance.

### Interindividual heterogeneity induces nonadditive changes in behaviour

Observed behaviour in the heterogeneous groups differed from the expectation based on the homogeneous groups (Fig. 3a). The direction and amount of such behavioural changes differed depending on the behavioural trait of interest and combinations of strains. Significant changes were found for spatial preference (*P* = 0.018) and stopping time (*P* = 0.018), indicating that the flies tended to prefer the centre of the arena and stop more frequently in the heterogeneous groups. There were no significant changes in moving speed (*P* = 0.8) and search comprehensiveness (*P* = 1.0).

**Figure 3.**
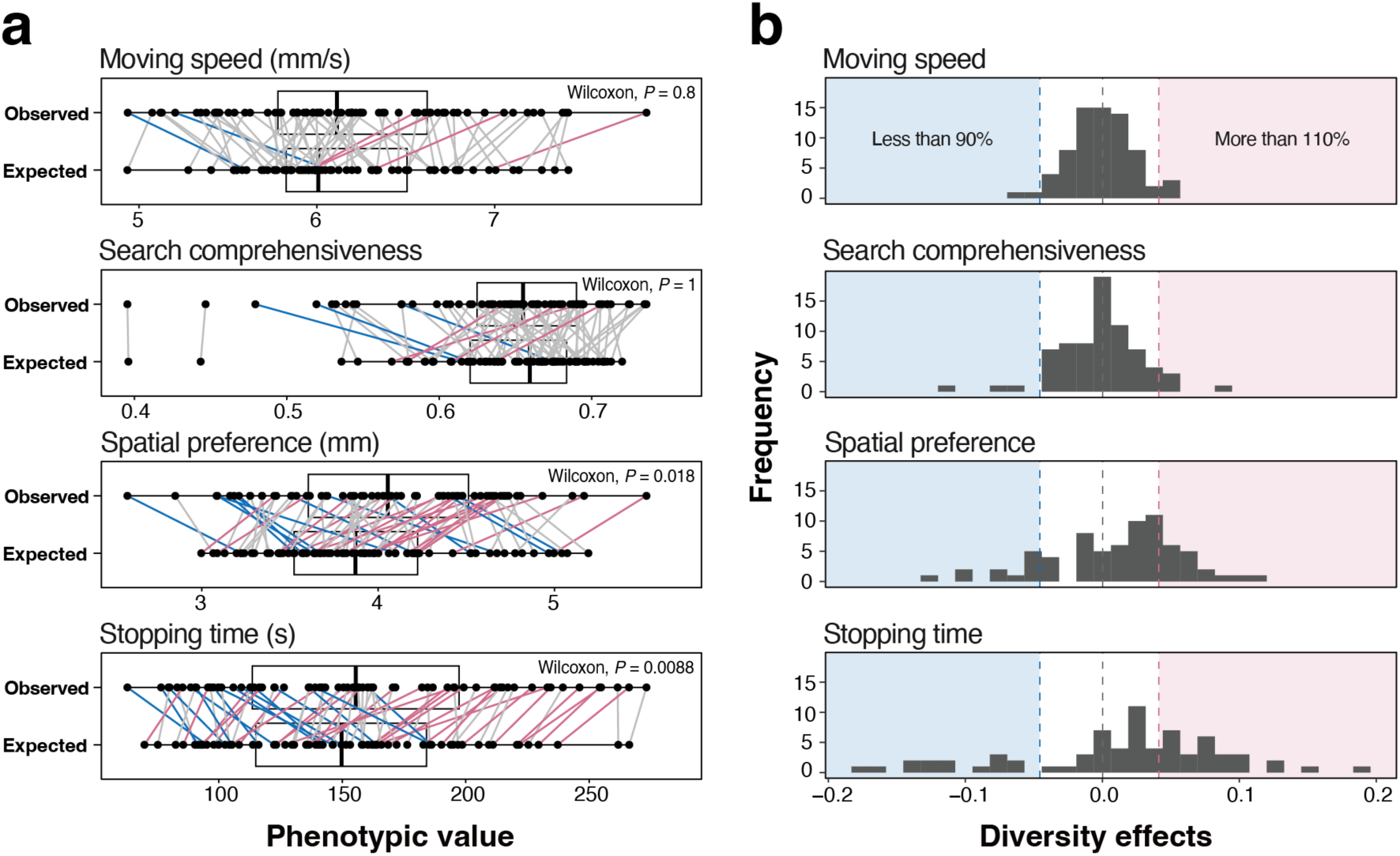
Behavioural changes due to the group composition. (a) Expected phenotypic values calculated from the homogeneous groups were compared with those observed in the heterogeneous groups. Combinations of strains with pink and blue lines indicate 10% increase and decrease in observed phenotypic values, respectively. Spatial preferences and stopping time significantly increased in the heterogeneous groups, as evaluated by Wilcoxon’s signed-rank test. (b) Diversity effects for each behavioural traits are shown in histograms, with coloured areas representing the amount of change. In all behavioural traits, the diversity effects varied widely across different strain combinations.

The calculated diversity effects showed wide variations among strain combinations and behavioural traits (Fig. 3b). Some combinations of strains demonstrated large diversity effects, and significant effects were specifically observed for stopping time.

### Behavioural foundations were found that explained the diversity effect

The results of PCA using the four behavioural indices revealed two major axes with high contribution (>25%), which summarised the behavioural characteristics. The PC1 axis was interpreted as the locomotor activity level, and the PC2 axis was associated with the exploratory range. A higher PC1 score represented the distance a fly walked in the 5 min, whereas a higher PC2 score indicated that the fly explored the arena more comprehensively. The summed counts of walking areas of representative strains with top or bottom five PC scores also supported this interpretation (Fig. 4a).

**Figure 4.**
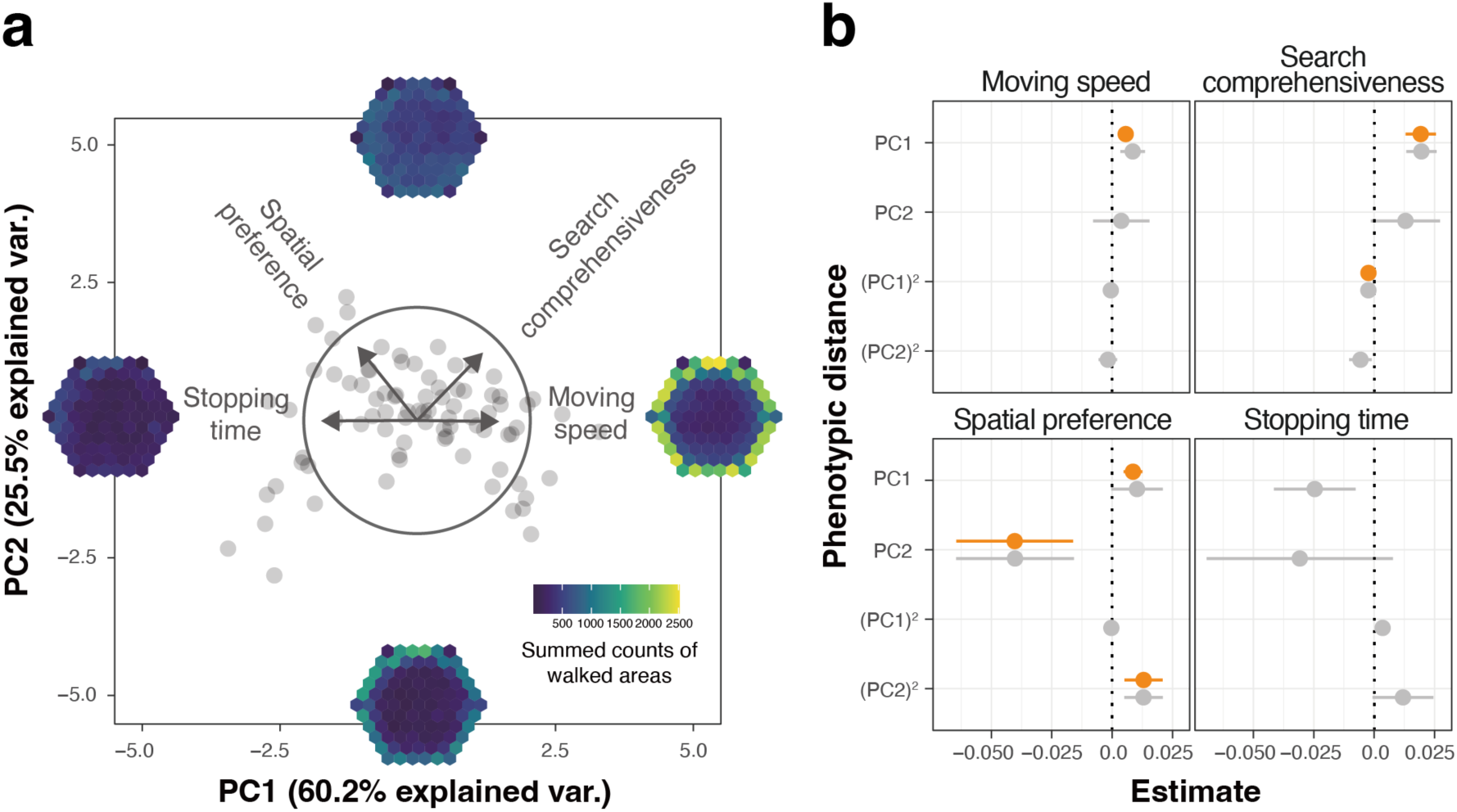
Phenotypic distances among strains possibly explain the diversity effect. (a) The results of principal component analysis with PC1 score on the horizontal axis and PC2 score on the vertical axis. The heatmaps show the summed counts of walked areas of the strains with the top or bottom five PC scores. (b) The results of multiple regression analysis are shown for the four behavioural traits. Orange and grey represent the results of the best and full models, respectively.

Using multiple regression analysis using the pairwise distance of PC1 and PC2 scores, we found that behavioural variations between strains can explain the diversity effects in the four behavioural traits (Fig. 4b). For instance, the distance of PC1 scores positively correlated with diversity effects in moving speed (*P* = 0.003), spatial preference (*P* = 0.030) and search comprehensiveness (*P* = 0.003), suggesting that greater disparities in the locomotor activity level of the two strains resulted in more pronounced positive changes in these behaviours. Moreover, the PC1 square term significantly predicted diversity effects in search comprehensiveness with a negative correlation (*P* = 0.039), indicating that when the difference of the locomotor activity level of the two strains was intermediate, the diversity effect of search comprehensiveness became higher. Finally, the diversity effect of spatial preference might be explained by the distance of PC2 scores with a negative correlation (*P* = 0.1) and its square term with a positive correlation (*P* = 0.1). These data suggested that when the exploratory range of the two strains was intermediate, the diversity effect of spatial preference became lower.

## Discussion

Although animal behaviour is primarily affected by innate genetic factors, behaviour can also be affected by conspecifics within a group (Frost et al., 2006). ‘Social facilitation’ is a phenomenon in which individuals from a group exhibit behavioural changes due to the presence of other individuals of the same species (Fonseca and Santos, 2023); this phenomenon has been reported in various flocking animals, including humans (Dindo et al., 2009), birds (Jackson et al., 2008), fish (Rosenthal et al., 2015) and fruit flies (Muria et al., 2020). The behavioural changes induced by behaviour of other members are known as the indirect genetic effect. The details of how the direction and strength of indirect genetic effects on group behaviour varied across different combinations of behavioural traits remain unclear. In this study, using a wide range of combinations from 83 genetically distinct strains, we explored the type and strength of heterogeneity leading to indirect genetic effects and synergistic effects on group behaviour. We detected positive synergistic changes in spatial preference and stopping time when two strains were combined, suggesting that behavioural heterogeneity among individuals induced social facilitation. Although most combinations of strains demonstrated no significant diversity effects, positive and negative diversity effects were observed for all four behavioural traits. This result indicated that the direction and strength of diversity effects varied with the composition of the group.

After exploring the group structure to explain the emergence of diversity effects, we found positive correlations between interindividual heterogeneity in the locomotor activity level (PC1) and the diversity effects of moving speed, search comprehensiveness and spatial preference. This result indicated that the presence of relatively active individuals affected conspecifics and altered group dynamics. A previous study showed that differences in the speed of locomotion among group members can result in the emergence of leaders (Gueron et al., 1996). Hence, differences in the level of locomotor activity may have induced the emergence of leaders and led individuals to synergistic behavioural changes. Conversely, it was interesting to find a negative correlation between interindividual heterogeneity in the exploratory range (PC2) and diversity effects of spatial preference, indicating that the spatial preference to the centre of the arena was enhanced by the presence of other individuals having a similar exploratory range. The spatial preference to the centre of the arena (risky, exposed area), so-called ‘boldness’, is a major behavioural trait often explored in studies on animal personalities. Boldness is a vital trait related to resource-seeking ability. A previous study on bass (*Perca fluviatilis*) demonstrated that shy individuals become bolder in groups composed of shy individuals than in groups consisting of bold individuals (Magnhagen and Staffan, 2005), which is partially consistent with our observations. Moreover, the diversity effects of search comprehensiveness and spatial preference negatively and positively correlated with the squared terms of PC1 (locomotor activity level) and PC2 (exploratory range) scores, respectively. These findings suggest that the effects of the strength of heterogeneity on the emergence of diversity effects are not always linear. Diversity effects can be maximised or minimised when the degree of heterogeneity is intermediate.

In the heterogeneous groups, stopping times were longer than expected. The behaviour of animals to stop moving is generally considered a behaviour to alert predators. Although the relationship between group size and vigilance behaviour towards predators has been extensively investigated (Beauchamp et al., 2021), our study is the first to demonstrate that heterogeneity within a group improves vigilance behaviour. Nevertheless, there is still no information on the group structure that causes the increase in stopping time, suggesting that the behavioural traits investigated in this study are limited towards fully explaining this phenomenon. Therefore, to understand the mechanism underlying the emergence of the diversity effect, it is necessary to quantify heterogeneity using a broader range of behavioural traits and conduct an exhaustive analysis.

This study describes the effects of interindividual heterogeneity on the dynamics of group behaviour; however, it is still our limitation that we could not determine whether and how the increase or decrease in behavioural traits influences the reproductive success of individuals. Hence, to understand the significance of aggregation with heterogeneous individuals, it is necessary to explore the impact of behavioural changes caused by heterogeneity on foraging success as well as reproductive success.

Altogether, our study has elucidated previously neglected by-productive effects of genetic heterogeneity on group dynamics in *Drosophila*. To our knowledge, this is the first study to demonstrate that heterogeneity among individuals in certain behavioural traits resulted in nonadditive behavioural modifications within groups. In addition to the conventional understanding of the effects of genetic diversity, where heterogeneity confers robustness to populations against environmental changes, our study provides a novel perspective on the effects of genetic diversity based on behaviour, i.e. heterogeneity can induce changes in behaviour through individual interactions and enable the acquisition of novel traits that would not otherwise be achievable. We suggest that such diversity effects should be considered a benefit of aggregation. Understanding the function of diversity effects in detail would help elucidate the significance of aggregation.

## Acknowledgments

This work was supported by the Japan Society for the Promotion of Science (Grants-in-Aid for Scientific Research JP23H03840 to Y.T.).

## Author contributions

T.O., D.X.S. and Y.T. conceived and designed the study. T.O. collected and analysed behavioural data of fruit flies with the help of D.X.S. and Y.T. Y.T. acquired the funding. T.O. wrote the first draft of the manuscript and all authors approved the final manuscript.

## Data availability

10.6084/m9.figshare.26242142 (for review: https://figshare.com/s/ce7b941c8b4de93ef48a)

---

**Figure S1.**
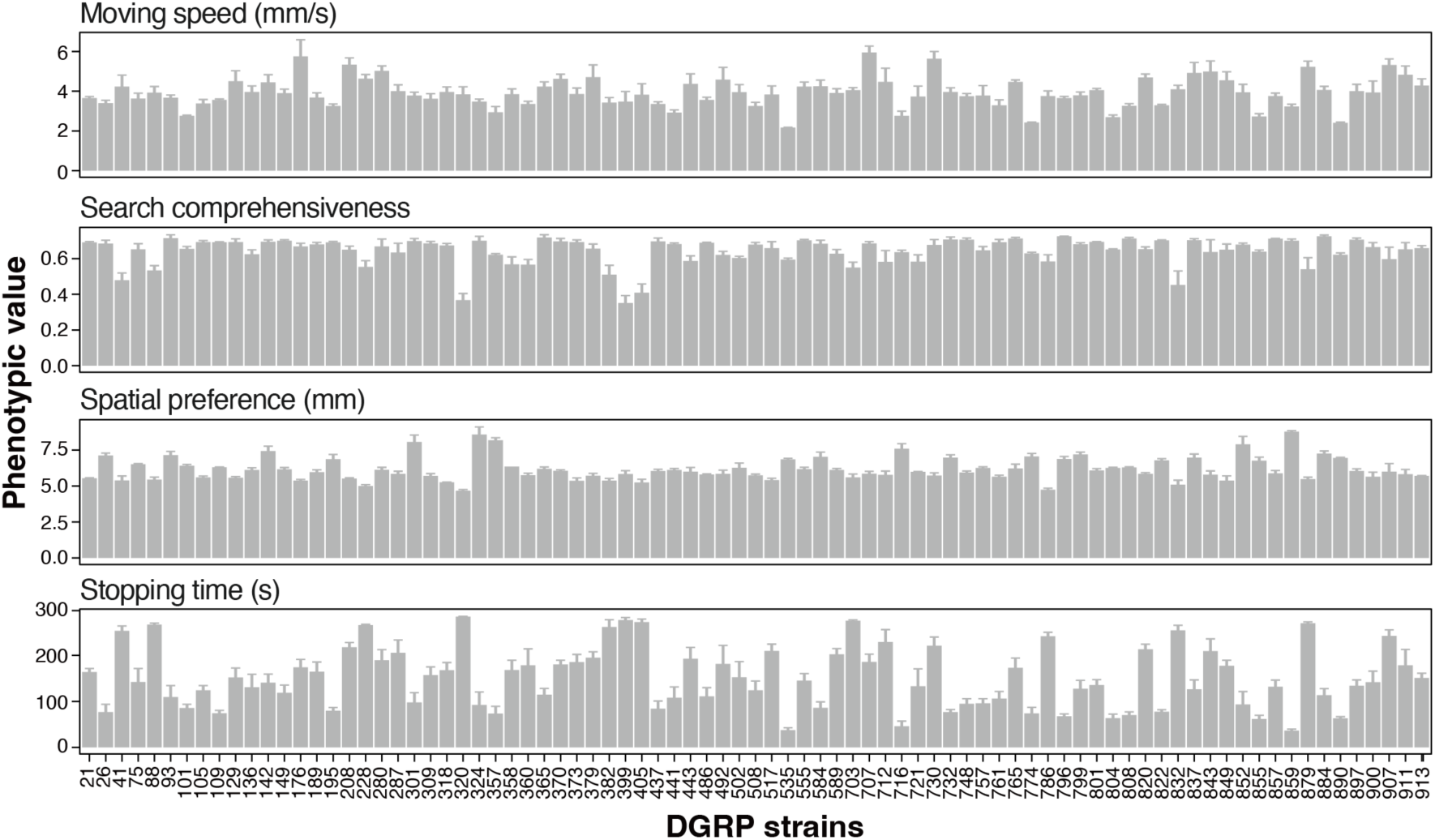
Four behavioural traits measured for the 83 strains used. Error bars represent standard errors of mean.

**Figure S2.**
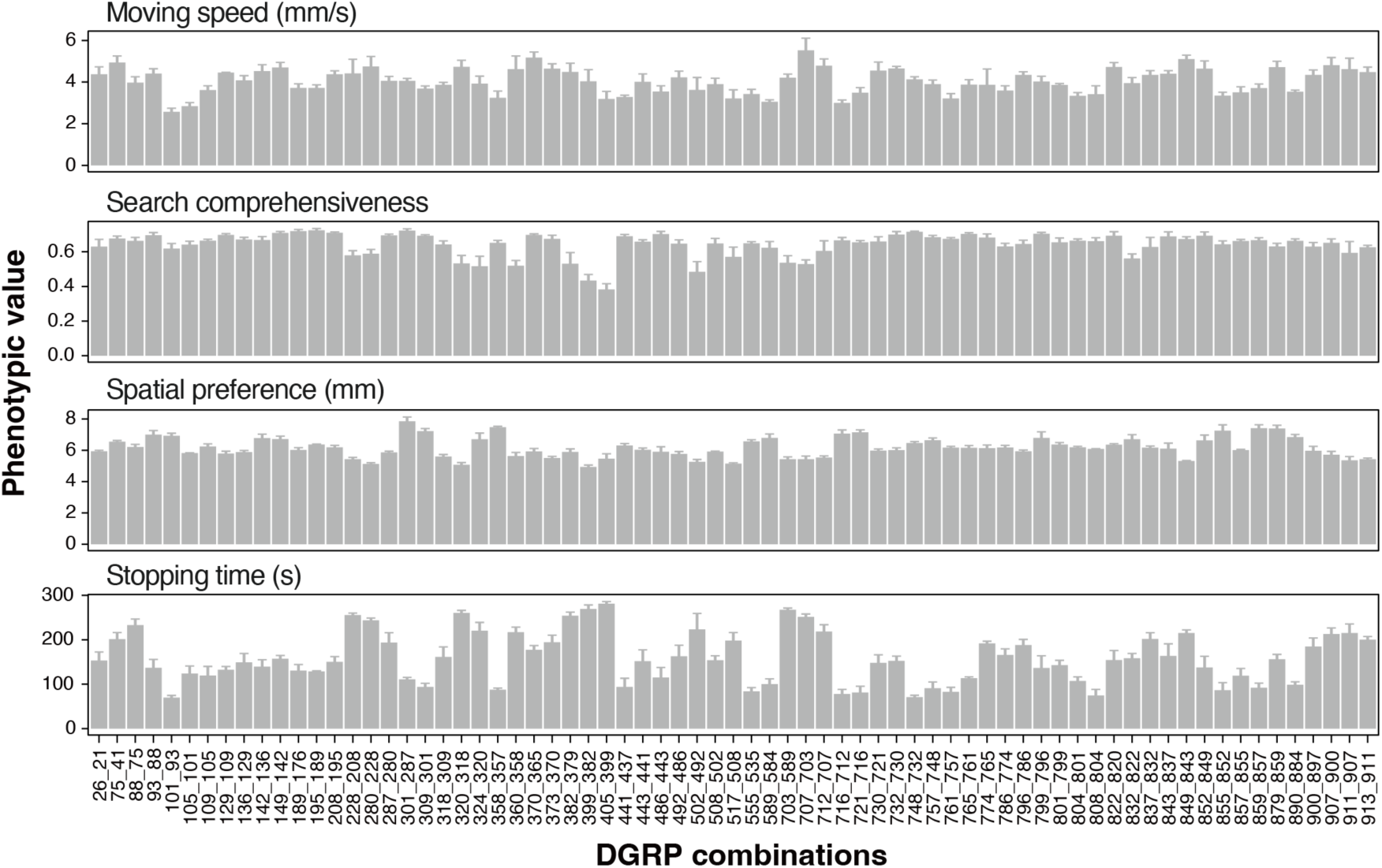
Four behavioural traits measured for the 71 combinations of fly strains used. Error bars represent standard errors of mean.

## References

1. Barnard, C. J. and Sibly, R. M. (1981). Producers and scroungers: A general model and its application to captive flocks of house sparrows. Anim. Behav. 29, 543–550.

2. Beauchamp, G., Li, Z., Yu, C., Bednekoff, P. A. and Blumstein, D. T. (2021). A meta-analysis of the group-size effect on vigilance in mammals. Behav. Ecol. 32, 919–925.

3. Biro, D., Sumpter, D. J. T., Meade, J. and Guilford, T. (2006). From compromise to leadership in pigeon homing. Curr. Biol. 16, 2123–2128.

4. Brandl, H. B., Pruessner, J. C. and Farine, D. R. (2022). The social transmission of stress in animal collectives. Proc. R. Soc. B Biol. Sci. 289, 20212158.

5. Colinet, H. and Renault, D. (2012). Metabolic effects of CO_2_ anaesthesia in *Drosophila melanogaster*. Biol. Lett. 8, 1050–1054.

6. Dindo, M., Whiten, A. and de Waal, F. B. M. (2009). Social facilitation of exploratory foraging behavior in capuchin monkeys (*Cebus apella*). Am. J. Primatol. 71, 419–426.

7. Dyer, J. R. G., Croft, D. P., Morrell, L. J. and Krause, J. (2009). Shoal composition determines foraging success in the guppy. Behav. Ecol. 20, 165–171.

8. Eyjolfsdottir, E., Branson, S., Burgos-Artizzu, X. P., Hoopfer, E. D., Schor, J., Anderson, D. J. and Perona, P. (2014). Detecting social actions of fruit flies. In Computer Vision – ECCV 2014 (ed. Fleet, D.), Pajdla, T.), Schiele, B.), and Tuytelaars, T.), pp. 772–787. Cham: Springer International Publishing.

9. Fonseca, A. and Santos, C. (2023). Reviewing social facilitation in insects over the past 30 years. Sociobiology 70, e9210.

10. Frost, A. J., Winrow-Giffen, A., Ashley, P. J. and Sneddon, L. U. (2006). Plasticity in animal personality traits: does prior experience alter the degree of boldness? Proc. R. Soc. B Biol. Sci. 274, 333–339.

11. Giraldeau, L.-A., Beauchamp, G., Giraldeau, L.-A., Beauchamp, G., Giraldeau, L.-A. and Beauchamp, G. (1999). Food exploitation: Searching for the optimal joining policy. Trends Ecol. Evol. 14, 102–106.

12. Gueron, S., Levin, S. A. and Rubenstein, D. I. (1996). The dynamics of herds: from individuals to aggregations. J. Theor. Biol. 182, 85–98.

13. Hamilton, W. D. (1971). Geometry for the selfish herd. J. Theor. Biol. 31, 295–311.

14. Jackson, A. L., Ruxton, G. D. and Houston, D. C. (2008). The effect of social facilitation on foraging success in vultures: a modelling study. Biol. Lett. 4, 311–313.

15. Jolles, J. W., King, A. J. and Killen, S. S. (2020). The role of individual heterogeneity in collective animal behaviour. Trends Ecol. Evol. 35, 278–291.

16. Keiser, C. N., Pinter-Wollman, N., Ziemba, M. J., Kothamasu, K. S. and Pruitt, J. N. (2018). The primary case is not enough: Variation among individuals, groups and social networks modify bacterial transmission dynamics. J. Anim. Ecol. 87, 369–378.

17. Kenward, R. E. (1978). Hawks and doves: Factors affecting success and selection in goshawk attacks on woodpigeons. J. Anim. Ecol. 47, 449–460.

18. Krause, J., Ruxton, G. D., Krause, J. and Ruxton, G. D. (2002). Living in Groups. Oxford, New York: Oxford University Press.

19. Mackay, T. F. C., Richards, S., Stone, E. A., Barbadilla, A., Ayroles, J. F., Zhu, D., Casillas, S., Han, Y., Magwire, M. M., Cridland, J. M., et al. (2012). The *Drosophila melanogaster* genetic reference panel. Nature 482, 173–178.

20. Magnhagen, C. and Staffan, F. (2005). Is boldness affected by group composition in young-of-the-year perch (*Perca fluviatilis*)? Behav. Ecol. Sociobiol. 57, 295–303.

21. Markham, A. C., Gesquiere, L. R., Alberts, S. C. and Altmann, J. (2015). Optimal group size in a highly social mammal. Proc. Natl. Acad. Sci. 112, 14882–14887.

22. Marzluff, J. M., Heinrich, B. and Marzluff, C. S. (1996). Raven roosts are mobile information centres. Anim. Behav. 51, 89–103.

23. Muria, A., Musso, P.-Y., Durrieu, M., Portugal, F. R., Ronsin, B., Gordon, M., Jeanson, R. and Isabel, G. (2020). Social facilitation of long-lasting memory is mediated by CO_2_ in *Drosophila*.

24. Neill, S. R. J. and Cullen, J. M. (1974). Experiments on whether schooling by their prey affects the hunting behaviour of cephalopods and fish predators. J. Zool. 172, 549– 569.

25. Perron, J. M., Huot, L., Corrivault, G.-W. and Chawla, S. S. (1972). Effects of carbon dioxide anaesthesia on *Drosophila melanogaster*. J. Insect Physiol. 18, 1869–1874.

26. Reynolds, C. W. (1987). Flocks, herds and schools: A distributed behavioral model. SIGGRAPH Comput Graph 21, 25–34.

27. Rosenthal, S. B., Twomey, C. R., Hartnett, A. T., Wu, H. S. and Couzin, I. D. (2015). Revealing the hidden networks of interaction in mobile animal groups allows prediction of complex behavioral contagion. Proc. Natl. Acad. Sci. 112, 4690–4695.

28. Santostefano, F., Wilson, A. J., Niemelä, P. T. and Dingemanse, N. J. (2017). Indirect genetic effects: a key component of the genetic architecture of behaviour. Sci. Rep. 7, 10235.

29. Simpson, E. H. (1949). Measurement of diversity. Nature 163, 688–688.

30. Takahashi, Y., Tanaka, R., Yamamoto, D., Noriyuki, S. and Kawata, M. (2018). Balanced genetic diversity improves population fitness. Proc. R. Soc. B Biol. Sci. 285, 20172045.

31. Treherne, J. E. and Foster, W. A. (1981). Group transmission of predator avoidance behaviour in a marine insect: The trafalgar effect. Anim. Behav. 29, 911–917.

32. Triplet, P., Stillman, R. A. and Goss-Custard, J. D. (1999). Prey abundance and the strength of interference in a foraging shorebird. J. Anim. Ecol. 68, 254–265.

33. Weimerskirch, H., Martin, J., Clerquin, Y., Alexandre, P. and Jiraskova, S. (2001). Energy saving in flight formation. Nature 413, 697–698.

34. Westneat, D., Fox, C., Westneat, D. and Fox, C. eds. (2010). Evolutionary Behavioral Ecology. Oxford, New York: Oxford University Press.

